# Experimental evolution of metabolism under nutrient restriction: enhanced amino acid catabolism and a key role of branched-chain amino acids

**DOI:** 10.1101/2023.02.06.527241

**Authors:** Fanny Cavigliasso, Loriane Savary, Jorge E. Spangenberg, Hector Gallart-Ayala, Julijana Ivanisevic, Tadeusz J. Kawecki

## Abstract

Periodic food shortage is a common ecological stressor for animals, likely to drive physiological and metabolic adaptations to alleviate its consequences, particularly for juveniles that have no option but to continue to grow and develop despite undernutrition. Here we study changes in metabolism associated with adaptation to nutrient shortage, evolved by replicate *Drosophila melanogaster* populations maintained on a nutrient-poor larval diet for over 240 generations. In a factorial metabolomics experiment we showed that both phenotypic plasticity and genetically-based adaptation to the poor diet involved wide-ranging changes in metabolite abundance; however, the plastic response did not predict the evolutionary change. Compared to non-adapted larvae exposed to the poor diet for the first time, the adapted larvae showed lower levels of multiple free amino acids in their tissues – and yet they grew faster. By quantifying accumulation of the nitrogen stable isotope ^15^N we show that adaptation to the poor diet led to an increased use of amino acids for energy generation. This apparent “waste” of scarce amino acids likely results from the trade-off between acquisition of dietary amino acids and carbohydrates observed in these populations. The three branched-chain amino acids (leucine, isoleucine and valine) showed a unique pattern of depletion in adapted larvae raised on the poor diet. A diet supplementation experiment demonstrated that these amino acids are limiting for growth on the poor diet, suggesting that their low levels resulted from their expeditious use for protein synthesis. These results demonstrate that selection driven by nutrient shortage not only promotes improved acquisition of limiting nutrients, but also has wide-ranging effects on how the nutrients are used. They also show that the abundance of free amino acids in the tissues does not, in general, reflect the nutritional condition and growth potential of an animal.

**Lay summary:** Juvenile animals are particularly vulnerable to nutrient deprivation – they usually do not have an option of arresting their development and just trying to survive until food becomes plentiful; rather, they must attempt to grow and develop with whatever nutrients that can scrape. While they could obviously improve their lot by getting better at finding the scarce food, could they also adapt their physiology and metabolism in a way that would alleviate consequences of undernutrition? To find out we let populations of the fruit fly *Drosophila melanogaster* adapt during 240 generations to conditions of chronic larval nutrient shortage, and then studied their metabolism. We found that these populations evolved changes in their amino acid metabolism: their larvae are better at extracting amino acids from nutritionally poor diet and are able to grow faster (which requires a higher rate of protein synthesis), while maintaining lower levels of most amino acids in their system. This suggests improved cellular “logistics”, with a higher turnover of raw materials associated with their lower stocks owing to their more efficient and immediate use. However, paradoxically, the malnutrition-adapted larvae also “waste” a substantial fraction of their amino acids by “burning” them for energy. They can afford this because of their improved extraction of scarce amino acid from the diet, but they may also be compelled to do this by a trade-off with absorption of dietary carbohydrates.

## Introduction

Many animal populations experience periods of nutrient shortage that result in physiological stress and reduced fitness. If these periods occur repeatedly over generations, they will generate natural selection for physiological adaptations that alleviate their consequences. These adaptations include accumulating metabolic reserves when nutrients are plentiful, and the ability to shut down reproduction and switch to a “survival mode”. This mode is characterized by reduced metabolism, mobilization of fat stores and, in extremis, autophagy, when nutrients become scarce (Tatar *et al*. 2003; Singh *et al*. 2019). They help the animal wait out famines and survive to reproduce once food becomes plentiful again. However, these strategies are often unavailable to juveniles exposed to prolonged undernutrition. With some exceptions like the *dauer* larva of *Caenorhabditis elegans*, juveniles cannot arrest their development and wait out the lean times; their only option is to attempt to develop and grow using whatever nutrients they can obtain. This is likely to favor adaptations in nutrient acquisition, metabolism and nutrient use different from those favoring famine survival in adults. Exploring these adaptations may thus help understanding aspects of metabolism and physiology of juvenile animals. Furthermore, given increasing evidence that aspects of metabolism appear programmed for life, it may contribute to explaining aging (de Magalhaes 2012; Carlsson *et al*. 2021) and human vulnerability to metabolic disease (Prentice 2005; Roseboom *et al*. 2006).

In this study we explore changes in metabolism associated with genetically-based adaptation to juvenile (larval) undernutrition evolved in a long-term laboratory natural selection experiment with *Drosophila melanogaster*. The experiment involves six “Selected” populations that evolved for more than 240 generations on a poor (nutritionally diluted) larval diet and six “Control” populations maintained in parallel on a standard (sufficient, but not exceedingly nutrient rich) diet. The poor diet results in a drastic reduction of larval growth, such that non-adapted larvae take more than twice as long to develop (from egg to pupa) than on standard diet, and yet emerge at less than half the weight of flies raised on the standard diet (Kolss *et al*. 2009; Vijendravarma *et al*. 2011). In the course of experimental evolution, the Selected populations acquired the ability to grow and develop substantially faster on the poor diet compared to Control populations (Kolss *et al*. 2009; Erkosar *et al*. 2017; Cavigliasso *et al*. 2020). They also evolved a smaller critical size for metamorphosis initiation (Vijendravarma *et al*. 2012); as a consequence, they emerge as smaller adults than Controls despite their faster growth on poor diet (Vijendravarma *et al*. 2011; Cavigliasso *et al*. 2020).

The faster growth of Selected larvae on poor diet appears at least partly mediated by improved acquisition of limiting nutrients. Selected larvae assimilate amino acids from poor diet at a higher rate than Controls, but are less efficient in assimilating sucrose (Cavigliasso *et al*. 2020), consistent with changes in the expression of digestive enzymes (Erkosar *et al*. 2017). An adaptive character of this change is supported by the finding that larval growth on the poor diet is limited by yeast (the main source of dietary protein) but not by sugar availability (Cavigliasso *et al*. 2020). Thus, experimental adaptation of Selected populations to larval undernutrition involved post-ingestive nutritional compensation, i.e., a shift in investment in the acquisition of the major categories of dietary macronutrients (Behmer 2009; Simpson *et al*. 2012).

However, it is unlikely that selection imposed by the dietary regimes was limited to mechanisms of nutrient acquisition. Whole genome sequencing identified signatures of allele frequency differentiation between Control and Selected populations at single nucleotide polymorphisms (SNPs) in over 100 genomic regions (Kawecki *et al*. 2021). These results point to a highly polygenic nature of adaptation to diet in these populations, suggesting the involvement of many physiological mechanisms. This observation is also supported by a large number of differentially expressed genes with diverse functions (Erkosar *et al*. 2017). Both candidate SNP-associated and differentially expressed genes include many genes involved in metabolism (notably in lipid and carbohydrate metabolism) and in the regulation of response to nutrients and growth, such as insulin and FOXO signaling (Erkosar *et al*. 2017; Kawecki *et al*. 2021). Therefore, the adaptation of Selected populations to the poor diet was mediated not only by changes in the acquisition of scarce nutrients but also by changes in the way the acquired nutrients were used in metabolism and growth.

Here we investigate such evolved differences in metabolism between Selected and Control larvae, particularly focusing on amino acids. Under the simplest scenario, the faster assimilation of amino acids from the poor diet by Selected larvae could directly feed into an increased rate of protein synthesis, leading to faster growth without significant changes in other aspects of metabolism. However, in addition to being building blocks of proteins, dietary amino acids are substrates for the synthesis of many important molecules, such as nucleotides, catecholamine hormones, neurotransmitters and cofactors (Wu *et al*. 2014), and some play a role in signaling (Manière *et al*. 2020). Furthermore, amino acids may be deaminated and the carboxylic skeleton may be used to generate ATP or energy stores (i.e., glycogen and fat) (Soultoukis & Partridge 2016; Holtof *et al*. 2019). This last use of amino acids by larvae faced with nutrient shortage might appear wasteful, especially that protein to carbohydrate ratio in both the standard and the poor diet is lower than optimal for *Drosophila* larvae (Rodrigues *et al*. 2015; Cavigliasso *et al*. 2020). Yet, processes that mediate, from foraging, through digestion and transport of nutrients, to protein synthesis, require a lot of energy. Furthermore, larvae need to accumulate triglyceride reserves to fuel the energetically costly process of metamorphosis (Boggs 2009). Because of their faster growth on poor diet, the Selected larvae presumably need more energy per time unit than Controls; yet, they are less efficient in assimilating dietary carbohydrates (Cavigliasso *et al*. 2020). These arguments lead to contrasting predictions: either selection under the poor diet regime may have favored minimizing the use of scarce amino acids for energy generation, or it may have favored their greater use to compensate for lower carbohydrate assimilation and increased energy need.

We first address the metabolic consequences of adaptation to poor diet with high-coverage targeted metabolomics, comparing the patterns of abundance of core metabolites, in particular amino acids and their derivatives and compounds involved in energy metabolism, between Selected and Control larvae. We employ a factorial design, whereby the metabolome of larvae from both Selected and Control populations is assessed on both poor and standard diets. Because the base population from which these populations were originally derived had been maintained on the standard diet for several years before the onset of the evolution experiment, it should have already been adapted to the standard diet. We therefore interpret metabolic differences between Selected and Control larvae raised on poor diet as a result of evolutionary change driven by selection imposed on the Selected populations by the poor diet regime. Differences between Selected and Control larvae raised on standard diet would then represent correlated responses to that selection.

Additionally, a comparison of the metabolome of larvae of the same population raised on poor *versus* standard diet represents a phenotypically plastic response of that population to nutritional conditions. This comparison is of interest in its own right, given the proposed role of free amino acids as indicators of nutrient availability, relied on by TOR pathway to regulate cell growth and division (Antikainen *et al*. 2017). However, the effect of dietary limitation on the level of amino acids and other metabolites in the larvae does not seem to have been assessed; in adult flies nutritional restriction strongly affects amino acid abundance but the direction of change differs between amino acids and studies (Avanesov *et al*. 2014; Laye *et al*. 2015; Jin *et al*. 2020).

We also explore the relationship between phenotypic plasticity and genetically-based adaptation. It is sometimes claimed that evolutionary adaptation to novel environments primarily occurs by natural selection favoring genetic variants that magnify the initial phenotypically plastic responses and eventually lead to their constitutive expression (Pigliucci & Murren 2003; Paenke *et al*. 2007; Lande 2009). However, evidence from gene expression patterns tends to be more consistent with a negative correlation between plastic responses and evolutionary changes (Yampolsky *et al*. 2012; Ghalambor *et al*. 2015; Huang & Agrawal 2016; Josephs *et al*. 2021). Several authors proposed that a positive (respectively negative) correlation between plasticity and evolution in gene expression patterns implies that plasticity is predominantly adaptive (respectively maladaptive) (Ghalambor *et al*. 2015; Huang & Agrawal 2016; Josephs *et al*. 2021). Our results suggest that this interpretation may be too simplistic.

The results of metabolic profiling revealed interesting patterns of differentiation in the abundance of amino acids and purine compounds, leading to two further specific questions that we addressed with additional experiments. First, we used amino acid supplementation to test for the role of branched-chain amino acids in adaptation to the poor diet. Second, we used a stable isotope-based approach to compare the rate of amino acid deamination between Selected and Control larvae. We describe the rationale for these experiments in the Results section below.

## Methods

This is an abbreviated outline of the methods; a detailed version is available as Supplementary Methods.

### Experimental evolution

We used six Control and six Selected populations of *Drosophila melanogaster* originally derived from an outbred base population adapted to the lab and to the standard diet (Kolss *et al*. 2009). The Control populations have been maintained at a controlled larval density on our standard yeast-cornmeal-sugar larval diet medium (1.25 % yeast w/v). The six Selected populations have been maintained on the same density on a poor larval diet containing one-fourth of the amount of nutrients of the standard diet. After their emergence, adult flies of both selection regimes were transferred to standard diet. The experiments described in this paper were performed after 244 to 287 generations of experimental evolution.

To obtain larvae for each experiment, we let adult flies from each population lay eggs overnight. The desired number of eggs was then transferred onto the experimental food media and inoculated with fly feces to ensure the homogeneity of larval microbiota. Parents and grandparents of the assayed larvae were all raised on standard diet to minimize non-genetic parental effects.

### Multiple Pathway Targeted Metabolomics

We performed targeted metabolomics on whole early third instar larvae, targeting metabolites involved in multiple core energy pathways (glycolysis, pentose phosphate pathway, amino-acid metabolism, purine and pyrimidine metabolism, TCA cycle and OXOPHOS, beta-oxidation). We implemented a factorial design, with larvae from all six Selected and six Control populations (factor regime) raised on both standard and poor diets (factor diet). We collected three replicate samples from each population on each diet (arranged in batches). Some samples were filtered out, leaving a total of 65 in which we quantified 174 metabolites. Log-transformed metabolite abundance, normalized to larval protein content, was analyzed with a linear mixed model, with diet (poor *versus* standard), regime (Selected *versus* Control) and their interaction as fixed factors, and the replicate population and batch as random factors.

### BCAA supplementation experiment

Metabolome results suggested alternative hypotheses about the patterns of abundance of branched-chain amino acids (BCAA) that led to contrasting predictions about the consequences of supplementing the poor diet with those amino acids, as described in Results below. To test these predictions, we raised larvae of Selected and Control populations on the poor diet and on the poor diet supplemented with the three BCAA (valine, leucine, and isoleucine) in amounts that resulted in doubling of their concentrations contained in yeast in the diets. To verify that any effect is specific to BCAA and not a consequence of adding any amino acids, we included a third treatment, with the poor diet supplemented with three other essential amino acids, tryptophan, histidine, and lysine, also doubling their concentration in the poor diet (WHK treatment). We recorded the egg-to-adult survival probability, the developmental rate (inverse of developmental time) and female dry weight and the combined developmental time and weight to estimate larval growth rate. This experiment was done in three blocks. To analyze the results, we fitted a generalized linear mixed model (for egg-to-adult survival probability) or linear mixed models (for the other traits) with treatment, regime, and their interaction as fixed factors, and the replicate population and block as random factors.

### ^15^N accumulation experiment

Metabolome data suggested differences in the rate at which amino acids are deaminated and the nitrogen excreted as waste. To verify this, we exploited the fact that the amino acid deamination process has a slight preference for the common light isotope of nitrogen (^14^N). As a consequence, proteins in tissues become enriched in the heavy isotope of nitrogen (^15^N) to a degree that increases with rate of use of amino-acids deamination (Minagawa & Wada 1984; Peterson & Fry 1987; Gannes *et al*. 1997). Because ^15^N is naturally rare, in this experiment we used food media containing yeast enriched in ^15^N, which improved signal to noise ratio.

We raised larvae from egg on the ^15^N-enriched standard and poor diet and collected samples of 20-30 prepupae (not sexed) and 20-30 freshly emerged virgin females. Two samples per population, diet and life stage were collected in two blocks separated in time. ^15^N content (δ^15^N value in per mil relative to standard air nitrogen, ‰ N_2_-Air) was measured in each sample and normalized to ^15^N in the diet. Δ^15^N (δ^15^N_prepupae or adult_ – δ^15^N_diet_) was analyzed with a linear mixed model with regime, diet, stage (prepupae *versus* adult females) and their interactions as fixed factors, and the replicate population and block as random factors.

## Results

### Metabolite abundance differs between Selected and Control larvae especially when raised on poor diet

As expected, the diet treatment had a strong impact on metabolite abundances, with principal component analysis (PCA) clearly separating the larvae raised on poor and standard diet, mainly along the first principal component axis (Figure 1A; Wilks λ = 0.01, F_3,18_ = 625.1, *P* < 0.001, MANOVA on PC1-3, Supplementary Table S1). In turn, the second and third PCs differentiated the Selected and Control populations (Figure 1A; Wilks λ = 0.14, F_3,18_ = 35.8, *P* < 0.001, MANOVA on PC1-3, Supplementary Table S1). The Selected larvae were differentiated from Controls in the PC1-3 space irrespective of diet (MANOVA contrasts, t_20_ = 8.8, *P* < 0.001 on poor diet, t_20_ = 5.2, *P* < 0.001 on standard diet). However, this difference was more pronounced on poor diet as manifested as regime × diet interaction in MANOVA on PC1-3 (Wilks = 0.58, F_3,18_ = 4.4, *P* = 0.018).

**Figure 1.**
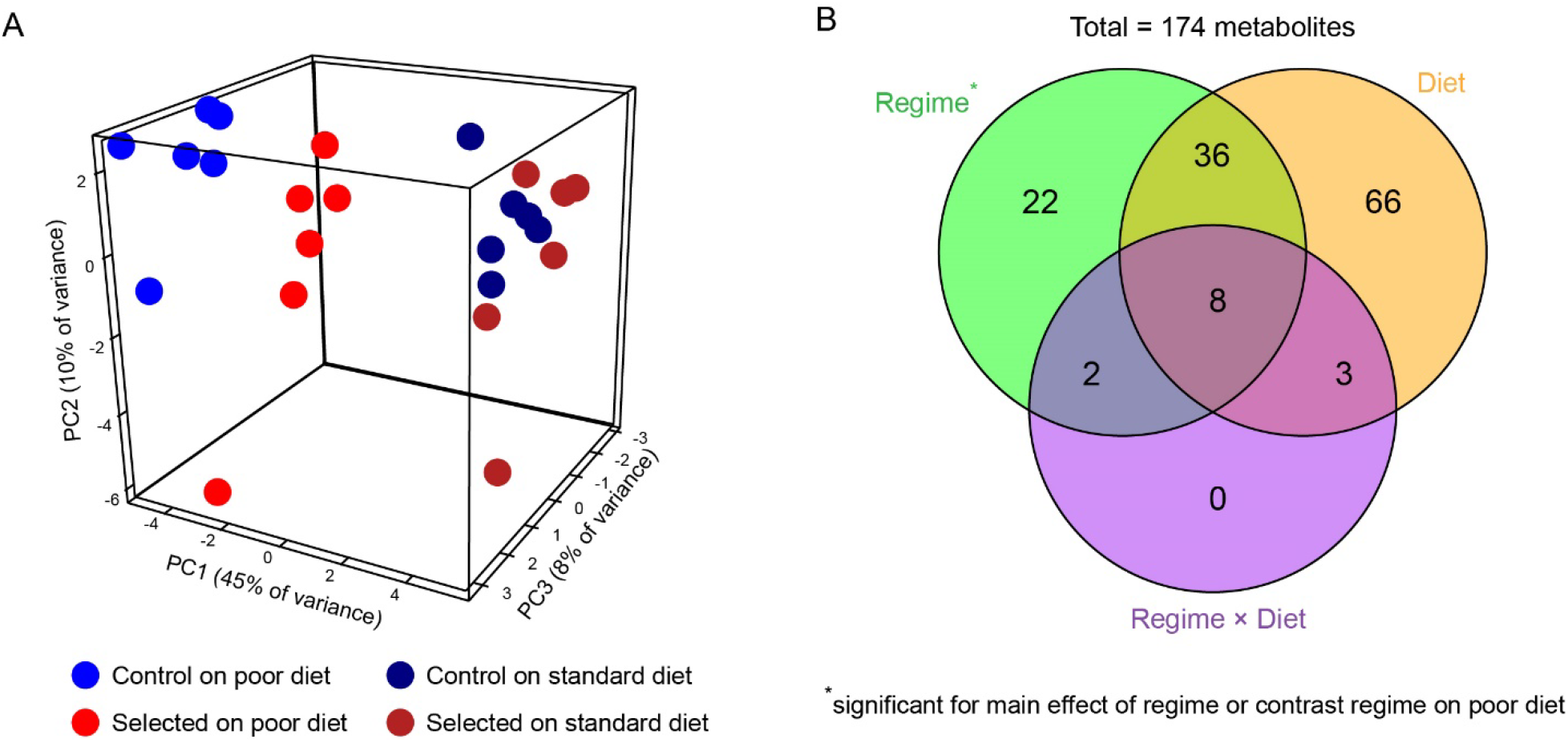
Diet quality induces both phenotypically plastic responses (“Diet”) and evolutionary changes (“Regime”) in metabolism. (A) PCA on the abundance of all 174 metabolites; each point corresponds to a population estimate averaged over replicate samples (an animated version is available in Supplementary Figure S1). B. Venn diagram summarizing the overlaps between significant metabolites. For diet this includes the main effect whereas for regime it includes both the main effect of regime and the regime contrast on poor diet (see text).

The median estimated effect size (absolute log_2_ fold-change) was twice as large for diet than for regime (0.28 *versus* 0.14, *P* < 0.001, Wilcoxon test). The median absolute effect of regime was smaller in larvae raised on poor than on standard diet (0.12 *versus* 0.19, *P* < 0.001, Wilcoxon test), but even the latter was still smaller than the effect of diet (0.19 *versus* 0.28, *P* < 0.001, Wilcoxon test). Thus, that the effect of diet on metabolome was generally larger than that of the evolutionary regime (Supplementary Figure S2).

Of 174 metabolites that passed quality assessment, univariate factorial analysis of abundance identified 113 affected by diet and 54 that showed a significant effect of regime (i.e., Selected *versus* Control populations) at 10% FDR. Although only 13 metabolites passed 10% FDR for regime × diet interaction, in a diet-specific contrast analysis 53 metabolites differed in abundance at 10% FDR between Control and Selected larvae raised on poor diet whereas only one differed on standard diet (Supplementary Table S2). In total 68 metabolites were detected as significantly different between Selected and Control lines at 10% FDR for as the main effect in the analysis (i.e., across diets) and/or in larvae raised on the poor diet. In the following we consider these 68 metabolites as significantly affected by evolution driven by diet (Figure 1B).

### Plastic response to poor diet involves accumulation of many metabolites

The response of each population to the diet treatment represents phenotypic plasticity (Scheiner 1993). Because the base population had been adapted to the standard diet and the Controls continued to experience the same diet during their evolution, the response of Control larvae to poor diet likely approximates the ancestral plastic response of the base population. This response would thus have been the starting point for evolutionary adaptation of the Selected populations when they were first exposed to the poor diet. We thus examined the effect of diet on metabolite abundance in Control larvae. We identified 44 (25%) metabolites as less abundant and 62 (36%) more abundant on the poor diet, spread among different metabolite groups (Figure 2, Supplementary Table S2). Metabolites with higher abundance on poor diet were particularly numerous among proteinogenic amino acids (11 out of 19), amino acid derivatives containing the amino group (24/50) and vitamins of B group (5/8). Thus, rather unexpectedly, dietary nutrient shortage resulted in an increase in tissue concentrations of many metabolites.

**Figure 2.**
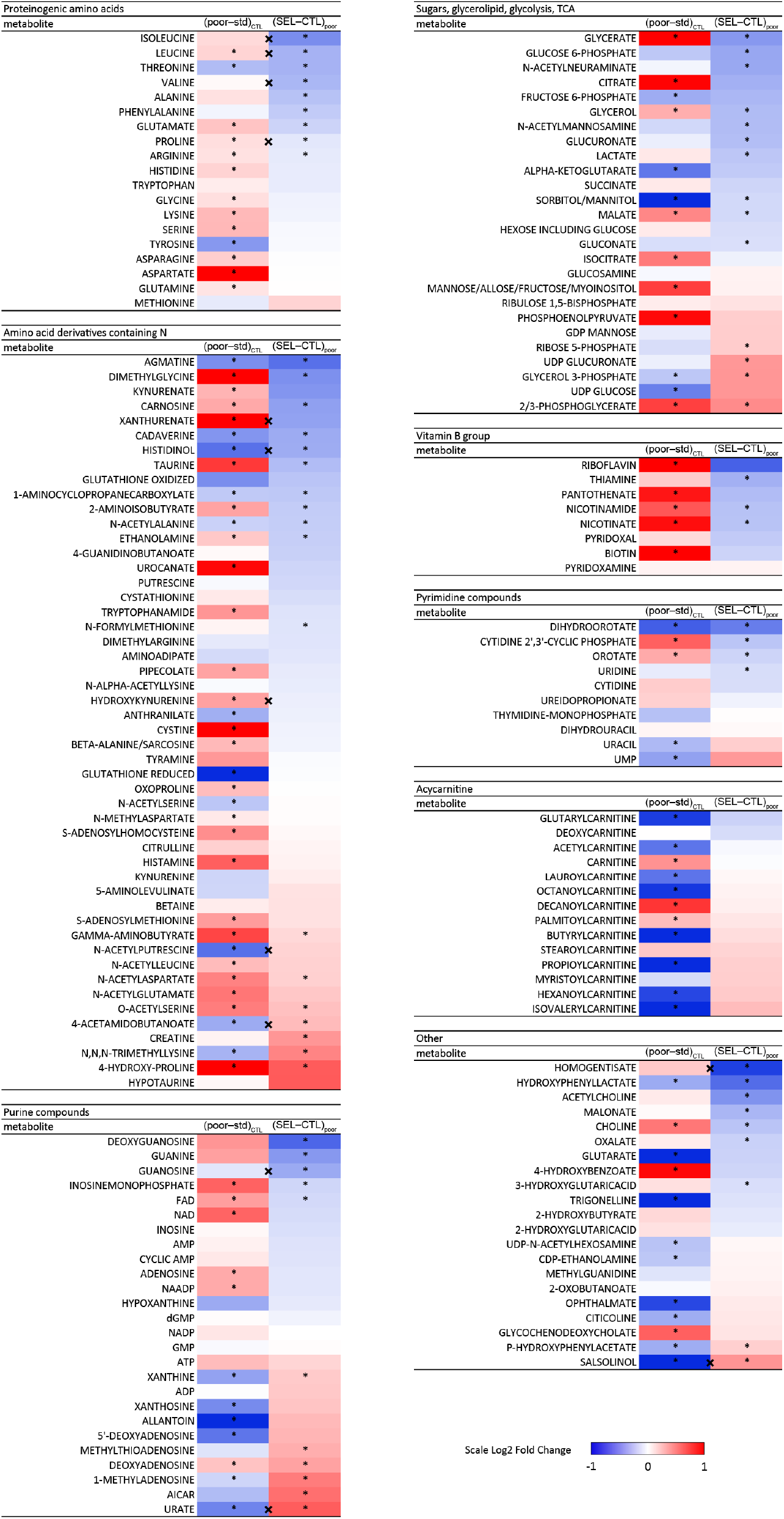
The effect of plastic response and evolutionary adaptation to poor larval diet on abundance of individual metabolites in 3rd instar larvae. For each metabolite, the first column of the heat map represents the estimated effect of diet (poor – standard) on Control larvae; the second column shows the effect of evolutionary regime (Selected – Control) on larvae raised on poor diet. \**q* < 0.1 for the corresponding contrast; (*) marks metabolites with *q* < 0.1 for the main effect of evolutionary regime but not for Selected – Control contrast on poor diet; “×” between columns indicates regime × diet interaction (*q* < 0.1, for patterns of those metabolites see Supplementary Figure S3). Several metabolites show diet effect with log-fold changes greater than ± 1 (Supplementary Figure S2); the color scale only varies between -1 and 1 to optimize resolution within this range. For a more traditional heatmap based on clustering of individual populations see Supplementary Figure S4.

### Adaptation to malnutrition involves changes in amino acid and purine metabolism

The phenotype expressed on the poor diet has been the target of selection in Selected populations. Differences between poor diet-raised Selected and Control thus represents the genetically-based response to selection evolved in the course of the 264 generations of experimental evolution driven by the diet. This evolutionary response is represented by the second columns of the heat maps in Figure 2. The 68 metabolites differentially abundant between Selected and Control larvae included chemically and functionally diverse compounds involved in a range of metabolic pathways. Nearly 3/4 of them (49) were less abundant in Selected than Control larvae while only 19 were more abundant, a significant deviation from 50:50 (sign test, *P* < 0.001). This includes lower abundance of several metabolites involved in sugar metabolism in Selected larvae, but higher abundance of a few that link glycolysis with triglyceride (2/3-phosphoglycerate, glycerol-3-phosphate) and glycogen (UDP-glucoronate) storage. Of the four building blocks of RNA, guanosine, and uridine were significantly less abundant while adenosine and cytidine tended to be less abundant. A rather consistent pattern was also apparent for amino acids; of 19 proteogenic amino acids, nine were significantly less abundant in Selected than Control populations while none was more abundant (Figure 2). Multiple amino acid derivatives containing the amino group were likewise affected by evolutionary regime, suggesting differences in amino acid catabolism. We also observed several changes in the purine / uric acid synthesis pathway, notably a higher concentration of AICAR, xanthine and urate in Selected compared to Control larvae. As we argue below, these differences suggest that Selected larvae catabolize a greater fraction of their amino acids, a hypothesis we test with an independent experiment.

### Plasticity of metabolite abundance in response to diet does not predict evolutionary response

As summarized in Introduction, plasticity is often proposed to anticipate evolutionary change driven by the same environmental factor. However, visual inspection of Figure 2 suggests no consistent relationship between the (presumably ancestral) plastic response of Controls to poor diet and the evolutionary change associated with genetically-based adaptation to it. For example, plasticity induced strong changes in abundance of acyl-carnitines (molecules transporting fatty acids in and out of mitochondria), but no such changes are detected between Selected and Control populations. Amino acids and vitamins tended to be more abundant on poor than standard diet but less abundant in Selected than Control larvae. Among purine compounds a few showed the same direction of change, but for many the evolutionary change tended in the direction opposite to the plastic response.

As another way of exploring the relationship between the evolutionary and plastic responses to poor diet, we plotted the plastic response of the Controls against the difference between Selected on poor and Controls on standard diet (Figure 3A). The latter reflects the difference in metabolome when each set of populations is raised under their evolutionary regime. Under one potential scenario (corresponding to Fig. 1B in Ghalambor *et al*. (2007)), the Control plastic response would represent a loss of homeostasis under a novel stress; evolution in Selected populations would then revert these changes, bringing the metabolome of Selected larvae on the poor diet to a similar state as that of Controls on standard diet. If so, most points in Figure 3A should have lain along the X-axis. This is clearly not the case; rather, most points lay along the diagonal and almost all metabolites that showed plastic response to diet have the same sign on X and Y-axes. Thus, even where evolution and plasticity acted in opposite direction, evolution usually only reduced the consequences of plasticity. Only for a few metabolites was the evolutionary change large enough to reverse the ancestral plastic response.

**Figure 3.**
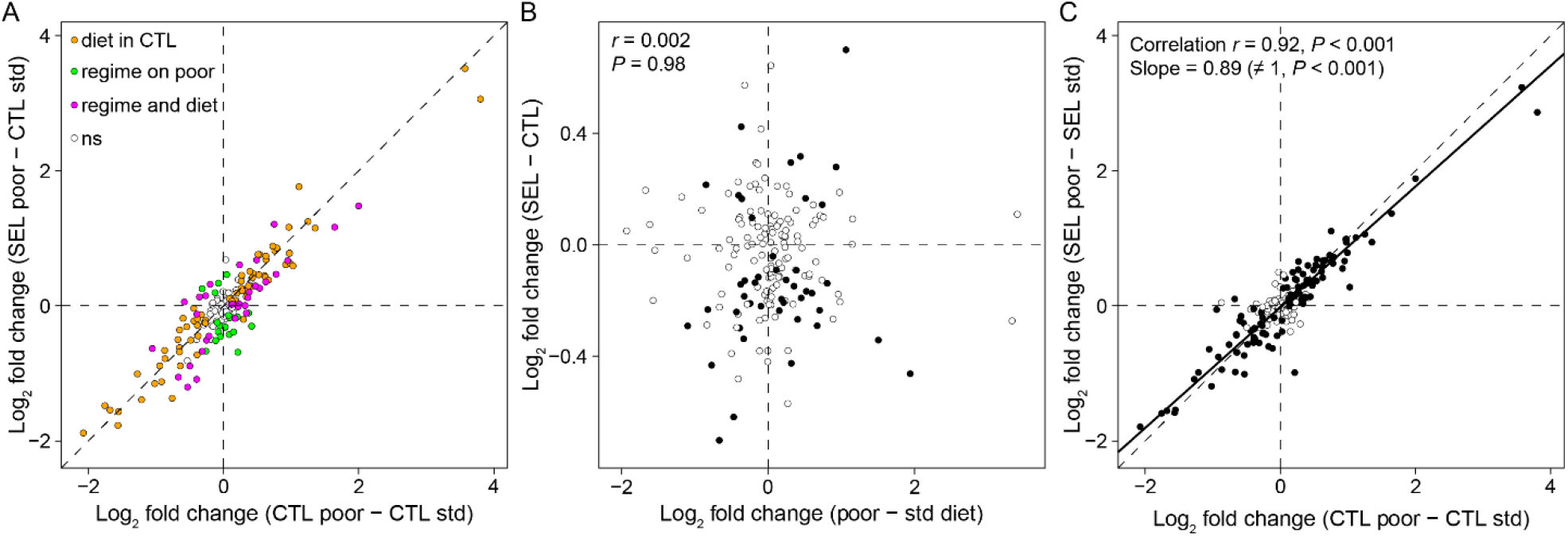
Plasticity *versus* evolutionary change. A. Adaptation to poor diet as modifier of ancestral plastic response: the relationship between metabolite abundance in Control and Selected larvae raised on poor diet, both expressed as a deviation from the ancestral state (Control larvae on poor diet). Colored metabolites are those with significant effect (*q* < 0.1) of diet in Controls (orange) or selection regime on poor diet (green) or both (magenta). CTL: Controls; SEL: Selected. Std: standard diet. B. The relationship between plastic and evolutionary responses to diet expressed as the main effects of diet and regime from the factorial model (see Methods). Each symbol corresponds to one metabolite, solid symbols are metabolites significant (*q* < 0.1) for the effects of both selection regime. C. The relationship between plastic responses of metabolome of Control and Selected populations. Solid symbols: metabolites significant (*q* < 0.1) for the main effect of diet; solid line represents the major axis regression fitted to all metabolites, dashed line is the diagonal. Correlations were calculated using all metabolites.

A quantitative assessment of the relationship between plasticity and evolution based on the values plotted in Figure 2 would be potentially biased due to non-independence of errors; therefore, we based such an assessment on the main effects of diet and regime from the factorial model (see Supplementary Methods). The overlap between metabolites significantly affected by diet and regime (44 shared metabolites; Figure 1B) was very close to the random expectation of 42.9 (Fisher’s Exact Test, *P* = 0.62). We also found no correlation between the estimated main effects of regime (Selected – Control) and diet (poor – standard), whether considering all metabolites (Pearson’s *r* = 0.002, *P* = 0.98) or only those significant for both effects (Figure 3B; *r* = 0.14, *P* = 0.36). In this last set, 21 metabolites showed the same sign of plasticity and evolutionary change (i.e., up in Selected and on poor diet or down in Selected and on poor diet) whereas 23 showed opposite signs, not different from random expectation (Fisher’s Exact Test, *P* = 1). These results imply that the plastic response of metabolome to diet did not generally predict the evolutionary response, whether in terms of metabolites affected or the direction of change.

We also found a signature of evolutionary change in the magnitude of plasticity. Even though the estimated effects of diet on metabolite abundance were highly correlated between the Control and Selected regimes, the major axis regression linking them had the slope of 0.89, slightly but significantly less than 1 (Figure 3C; *P* < 0.001). In other words, the response of Selected larvae to diet was on average 11% smaller in magnitude than that of Control larvae. This implies that metabolic state of Selected evolved to be somewhat more robust to diet change than that of Controls.

### BCAA are limiting for development on poor diet

We have previously shown that Selected larvae extract and absorb amino acids from poor diet at a faster rate than Controls (Cavigliasso *et al*. 2020). Thus, the lower concentration of many amino acids in Selected larvae may seem paradoxical. Among them, the three branched-chain amino acids (BCAA: valine, leucine, and isoleucine) show a unique pattern of regime × diet interaction. All BCAA are depleted in Selected larvae raised on the poor diet, but in Control larvae raised on poor diet they are at least as abundant as in either set of larvae raised on standard diet (Figure 4A-C). (Proline, the fourth amino acid with significant interaction is shows a different pattern; Figure 4D). One potential explanation is that maintaining a high level of free BCAA on poor diet is in itself maladaptive, e.g., because of their role in regulation of TOR signaling (Soultoukis & Partridge 2016; Juricic *et al*. 2020) or their correlation with development and physiological disturbance (Tsai *et al*. 2020). If so, Selected populations might have been selected to reduce their level. Under this scenario, supplementing the poor diet with BCAA would be expected to have no or negative effects on larval growth, development and/or survival. Alternatively, the lower level of free BCAA and several other amino acids in Selected larvae on the poor diet might reflect their more efficient or immediate use in protein synthesis or metabolism. In this scenario, the particularly strong reduction of BCAA might indicate that they are limiting for protein synthesis and thus growth and development on the poor diet; hence, supplementing the poor diet with BCAA should lead to improved larval performance.

**Figure 4.**
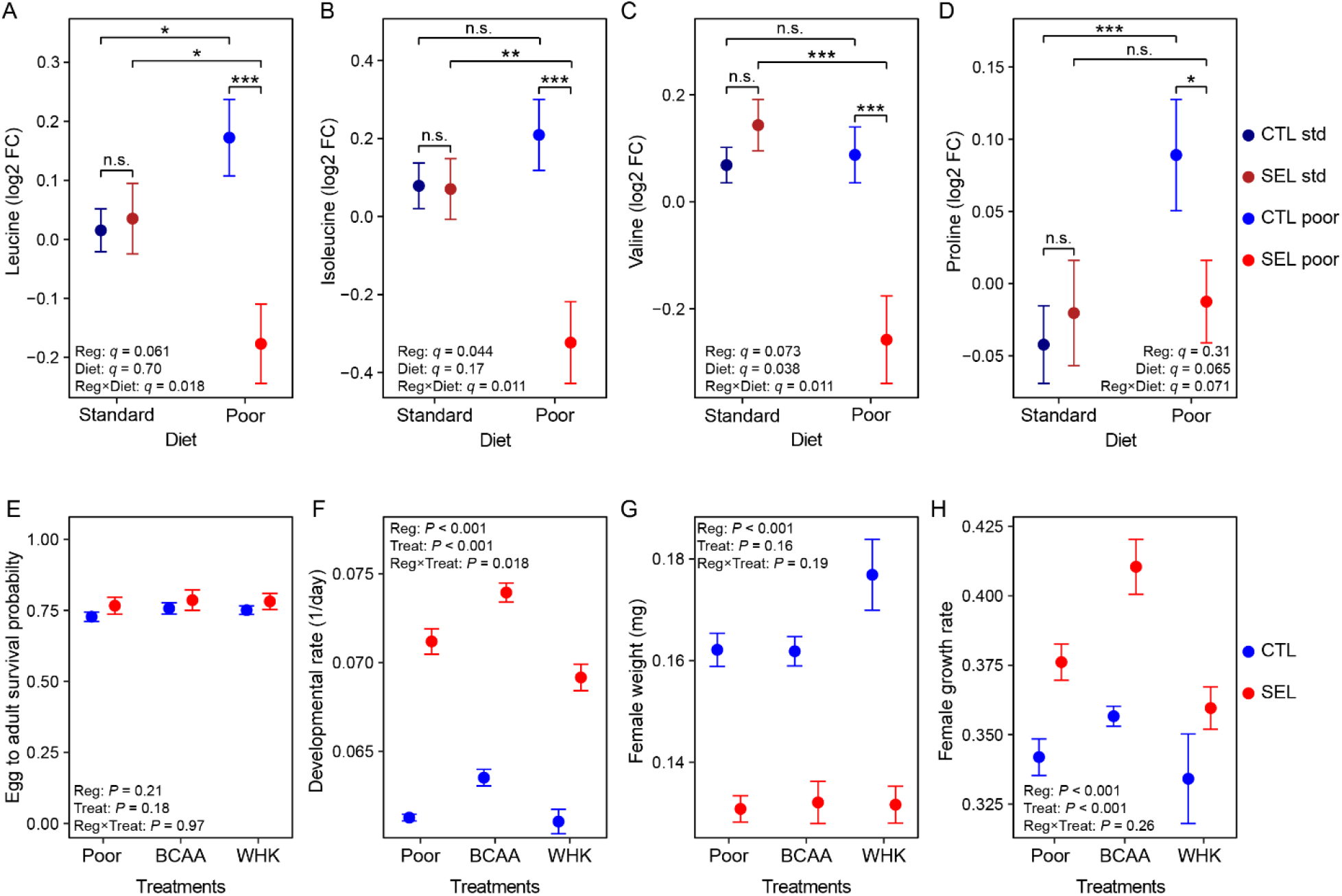
Branched-chain amino acids (BCAA) and adaptation to poor diet. Top row: Patterns of abundance (mean ± SE) of amino acids that show significant regime × diet interaction in the metabolome data: leucine (A), isoleucine (B), valine (C) (BCAA) and proline (D); n.s., *q* > 0.1; \**q* < 0.1; \*\**q* < 0.05; \*\*\**q* < 0.01. Bottom row: Effects of supplementing poor diet with BCAA or a mix of tryptophan, histidine and lysine (WHK) on egg-to-adult survival probability (E), developmental rate (F), female dry weight (G) and female estimated growth rate (H). Symbols indicate means ± SE. *N* = 3 replicate bottles per population and treatment. CTL: Controls. SEL: Selected. Reg: regime. Treat: treatment. Details of statistics in Supplementary Tables S3-S7.

To test these contrasting predictions, we studied how supplementation of poor diet with BCAA affects larval developmental traits in the Selected and Control populations. To verify that a potential effect is specific to BCAA rather than being mediated by any amino acids, we also investigated the effect of supplementation of the poor diet with a mix of three other essential amino acids, tryptophan, histidine, and lysine (referred to as WHK from one letter abbreviations of those amino acids) (Supplementary Figure S5).

Supplementing the poor diet with either BCAA or WHK did not change the egg-to-adult survival probability (Figure 4E; Supplementary Table S4). However, the developmental rate of Selected and Control populations did respond differently to amino acid supplementation (Figure 4F; Supplementary Table S5). This interaction was not driven by differential response to BCAA supplementation: adding BCAA improved developmental rate both Selected and Controls populations (t_21_ = 3.5, *P* = 0.006 for Controls, t_20_ = 4.6, *P* < 0.001 for Selected) and to a similar degree (interaction contrast (BCAA – Poor)_in SEL_ – (BCAA – Poor)_in CTL_: t_20_ = 0.7, *P* = 0.48). By contrast, supplementing poor diet with WHK tended to affect Control and Selected populations differently ((WHK – Poor)_in SEL_ – (WHK – Poor)_in CTL_: t_20_ = 2.1, *P* = 0.051), with no detectable change of developmental rate in Controls (WHK – Poor: t_20_ = 0.5, *P* = 0.90) and a negative effect in Selected larvae (Figure 4F; WHK – Poor: t_20_ = 3.4, *P* = 0.008).

We found no effect of amino acid supplementation on adult female dry weight (Figure 4G; Supplementary Table S6), despite the faster development of BCAA-supplemented larvae. Consistent with this, females raised on poor diet supplemented with BCAA had higher growth rate than those raised on poor diet (Figure 4H; BCAA – Poor: t_20_ = 2.8, *P* = 0.029; Supplementary Table S7). As reported before (Kolss *et al*. 2009; Erkosar *et al*. 2017; Cavigliasso *et al*. 2020), Selected larvae grew faster than Controls but still produced smaller adults because of shortened developmental time; however the effects of amino acid supplementation treatments on female weight and growth rate did not differ between Selected and Control populations (Supplementary Tables S6 and S7).

Thus, BCAA – and not the other three amino acids we tested – appeared to be limiting for larval growth and developmental rate on the poor diet, although they did not seem to affect the survival or adult weight.

### Selected larvae catabolize amino acids at a higher rate

The metabolome analysis revealed an apparent depletion of multiple amino acids in Selected larvae compared to Controls (Figure 2). In addition to a more immediate use of amino acids in protein synthesis, this could be mediated by increased use of amino acids for energy generation, a process that involves deamination and excretion of the resulting nitrogen waste. In contrast to mammals, in insects nitrogen liberated in amino acid catabolism enters the purine / uric acid synthesis pathway and is ultimately excreted as urate (uric acid) or its derivative allantoin (Salway 2018). Consistent with a higher deamination and nitrogen excretion rate, the Selected larvae show an increased concentration of urate and its immediate precursor xanthine, as well as of AICAR, a nucleotide that carries four nitrogen atoms from deamination into purine synthesis (Figure 2, Figure 5A-B).

**Figure 5.**
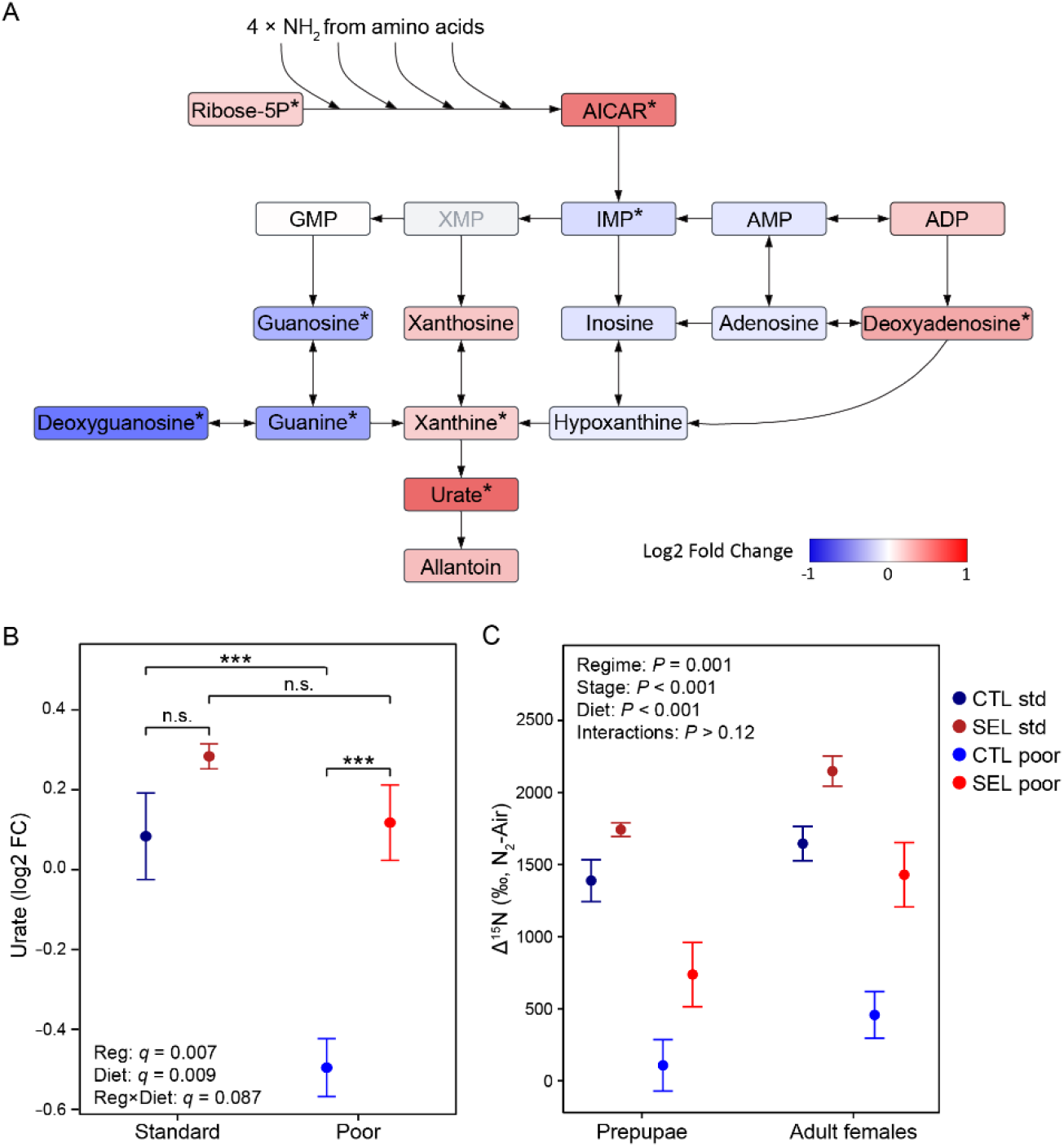
Plasticity and evolution of amino acid deamination in response to poor diet. A. Simplified purine/uric acid synthesis pathway (created in Lucidchart, www.lucidchart.com). Metabolites are colored according to their estimated fold change (Selected – Control) on poor diet; \**q* < 0.1. XMP was not reliably quantified. B. Relative concentration of urate in Selected and Control larvae raised on poor and standard diet. C. Isotope content after feeding on poor or standard diet enriched in ^15^N. Δ^15^N (‰ N_2_-Air) refers to the difference between δ^15^N of prepupae or adults and δ^15^N of the larval food. Symbols indicate means ± SE. Details of statistics in Supplementary Table S8. N = 2 samples of 20-30 individuals per population, diet and life stage.

To perform an independent test of this hypothesis we exploited the fact that proteins in tissues become enriched in ^15^N to a degree that increases with rate of use of amino-acids for energy generation (Minagawa & Wada 1984; Peterson & Fry 1987; Gannes *et al*. 1997). Thus, if Selected larvae have a higher deamination rate, they should accumulate more ^15^N than Control larvae raised on the same diet. Furthermore, if the level of urate and related compounds indeed is an indicator of the rate of amino acid deamination, larvae raised on the poor diet should accumulate less ^15^N than those raised on standard diet.

In agreement with our prediction, at the end of their larval development (at the prepupal stage), as well at eclosion, the Selected populations had a higher ^15^N content (i.e., higher Δ^15^N value) than Controls. This was the case on both poor and standard diet (Figure 5C; F_1,11_ = 18.9, *P* = 0.001; Supplementary Table S8). The Δ^15^N value of Control prepupae on poor diet was close to 0‰ (i.e., δ^15^N in prepupae was close to δ^15^N of the diet), suggesting negligible deamination (Figure 5C). Furthermore, individuals of both sets of populations accumulated more ^15^N when raised on standard *versus* poor diet (Figure 5C, F_1,12_ = 88.4, *P* < 0.001). Thus, consistent with the predictions derived from the metabolome analysis, larvae confronted with a nutrient-poor diet reduced the rate at which they catabolize amino acids, but the Selected larvae catabolized a greater fraction of assimilated amino acids than Controls.

## Discussion

### Plasticity, evolutionary adaptation, and metabolic efficiency

As expected, nutrient restriction imposed by the poor diet resulted in large changes in metabolite abundance. What was unexpected is that more metabolites increased than decreased in abundance in larvae raised on the poor diet relative to those raised on standard diet. The metabolome data are normalized to the sample protein content. Assuming that protein content is a measure of structural size of the larva, differences in normalized metabolite abundance should thus reflect differences in their relative concentration in the cells and hemolymph. Therefore, the greater abundance of core metabolites such as group B vitamins, nucleotide cofactors (FAD+, NAD+, NADP+, ATP) and multiple amino acids in larvae raised on the poor diet appears paradoxical: compared to the standard diet, the poor diet contains 25% of all dietary sources of these metabolites or their precursors. To our knowledge, no other study has reported consequences of larval dietary restriction for metabolome, but two studies report an increase in abundance of some amino acids in response to dietary restriction at an adult stage (Supplementary Table S5 in Laye *et al*. (2015), Supplementary Table A in Jin *et al*. (2020)). Thus, dietary availability of protein does not seem to lead to generally lower concentrations of amino acids in the tissues.

These changes in metabolite abundance obviously do not mean that the larvae are somehow better off on the poor diet. Possibly, the accumulation of some amino acids and most other metabolic changes in poor-diet raised larvae, at least those from the Control populations, reflect a stress-induced perturbation of homeostasis that hampers the optimal use of even the small quantity of nutrients the larvae manage to extract. Alternatively, the metabolic responses to poor diet might mostly reflect adaptive phenotypic plasticity in metabolism that alleviates the consequences of undernutrition. Even though the Control populations were not exposed to the poor diet during the experimental evolution, such adaptive plastic responses might have evolved in wild ancestors under variable nutrient availability in nature.

It has been argued that one could ascertain the adaptive *versus* non-adaptive (or maladaptive) nature of plasticity in response to a novel environmental stress by examining genetically-based evolutionary change driven by same stress. If the sign of plastic response and evolutionary change in a phenotype is the same (“synergistic”), it would suggest that the plastic response is adaptive; if the signs are opposite (“antagonistic”), the plastic response would be maladaptive (Yampolsky *et al*. 2012; Ghalambor *et al*. 2015; Huang & Agrawal 2016; Josephs *et al*. 2021). In this light, the studies just cited explored the correlation between phenotypically plastic and evolutionary changes in gene expression patterns, the first three finding a negative correlation and the last one a positive one. Here, we found no overall correlation between plastic and evolutionary responses of metabolite abundance to diet; for metabolites that showed a plastic response, evolution was as likely to act in the same as in the opposite direction to plasticity. However, nearly all metabolites that increased (decreased) in abundance in the Control larvae in response to the poor diet remained more (less) abundant in the Selected larvae on poor diet than in Control larvae on standard diet. In other words, even if for many metabolites the evolutionary change in metabolite abundance was antagonistic to plasticity, in most cases it was too small to completely offset or to reverse the plastic response. This can be interpreted in two ways. Possibly, in case of these metabolites the plastic response was maladaptive and thus opposed by selection in the Selected populations, but there was not enough time and/or genetic variability for evolution to bring the phenotype to the ancestral state. Alternatively, the initial plastic response may have been adaptive in terms of general direction, but imprecise, resulting in some degree of metabolic imbalance, with some metabolites overshooting the optimal level. Under the latter scenario, the phenotypic plasticity of metabolism may at the same time be adaptive (compared to no response) and induce some loss of metabolic homeostasis. If so, the evolutionary changes in either direction would represent fine-tuning of the initial plastic response.

We tend to favor the latter interpretation. A maladaptive plastic response would bring the phenotype farther away from the optimum than an adaptive response. Assuming that the strength of directional selection is greater farther away from the optimum, selection to revert maladaptive plastic responses should be stronger than selection to refine adaptive responses. Yet, we see no general tendency for the evolutionary changes that are antagonistic to the initial plasticity to be larger in magnitude than those that are synergistic with it, suggesting that plastic responses opposed by evolutionary change are not more maladaptive than those where evolution acts in the same direction as plasticity. While this is a speculation, this argument illustrates that it may be too simplistic to interpret plastic responses as adaptive or not solely based on the sign of correlation between plasticity and evolutionary change driven by the same environmental factor.

While the plastic response to poor diet led to more increases than decreases in metabolite abundance, of the metabolites affected by evolutionary adaptation to the poor diet nearly 3/4 decreased. This trend held regardless of the presence and direction of the plastic response of these metabolites. Many of these metabolites are building blocks of macromolecules (amino acids, nucleotides and nucleosides), precursors of cell membrane lipids and intermediates in energy metabolism. Some of them can be derived from dietary sugars and so their lower levels might be explained by the lower assimilation of carbohydrates by Selected than Control larvae (Cavigliasso *et al*. 2020). However, the Selected larvae assimilate amino acids from poor diet at a higher rate than Controls (Cavigliasso *et al*. 2020). Thus, rather than reflecting a supply bottleneck, the lower abundance of amino acids in Selected larvae may result from their more efficient use. The key metabolic cofactors NAD+, NADP+ and FAD+ and their vitamin precursor were also lower in Selected than Control larvae on the poor diet, but remained in excess of the level of Control larvae on standard diet and thus unlikely to be deficient to the degree that would limit metabolic flux. While we do not have any direct measure of metabolic fluxes, the fact that on poor diet Selected larvae grow faster than Controls implies that they must achieve a higher rate of protein, nucleic acid and membrane lipid synthesis, as well as being able to provide for the energy cost of the faster growth. Indeed, despite lower abundance of many energy metabolism intermediates and electron-carrying cofactors, Selected larvae do not have a lower level of ATP than Controls. Our results thus support the notion that experimental adaptation to larval undernutrition has been mediated in part by more efficient metabolism that reduced the concentration of cellular “building blocks” and metabolic intermediates while supporting faster growth in the face of dietary nutrient limitation. They also show that the abundance of free amino acids does not in general reflect the nutritional condition of an animal and its growth potential.

### Trade-off in nutrient acquisition leads to “waste” of amino acids for energy generation

Among the minority of metabolites whose abundance on poor diet was higher in Selected than Control larvae, most are amino acid derivatives. While some have known physiological functions in the nervous system (GABA, N-acetylaspartate) or muscles (creatine), many are products of amino acid degradation with no known physiological function, including the main nitrogenous excretion compound urate (uric acid). Consistent with the pattern of urate abundance, our ^15^N accumulation experiment demonstrated that, while Control larvae nearly completely cease deaminating amino acids on the poor diet, in the Selected larvae the rate of deamination on poor diet was restored to about half of that observed on standard diet. This implies that the ancestral plastic response to nutrient shortage was to minimize “wasting” of amino acids, but that adaptation to the poor diet favored a moderate use amino acids for energy generation.

This might be explained by the Selected populations having evolved the ability to assimilate amino acids from the poor diet at a higher rate (Cavigliasso *et al*. 2020). However, Selected larvae still grow considerably more slowly on the poor than standard diet. Arguably, using these amino acids for protein synthesis instead of deaminating them might enable an even faster growth and development. Faster growth requires increased supply of ATP; just protein synthesis consumes about 5 ATP molecules per attached amino acid (Marchingo & Cantrell 2022). Yet, the Selected larvae have become less efficient in extracting carbohydrates from the poor diet (Cavigliasso *et al*. 2020). While some of this reduction in carbohydrate acquisition may be compensated by lower accumulation of fat stores (Cavigliasso *et al*. 2020), it may also lead to a gap between the energetic needs and the supply of carbohydrate energy sources. In other words, it is not just that the Selected larvae and pupae can afford to “waste” some amino acids for energy; they may be compelled to do so by an evolutionary trade-off in acquisition of amino acids *versus* carbohydrates from the diet.

### A special role for branched chain amino acids in adaptation to undernutrition?

The three branched-chain amino acids – leucine, isoleucine and valine – showed a unique pattern of interaction, remaining similar or increasing in response to poor diet in Control larvae but strongly declining in Selected larvae (Fig 4 A-C). This was mediated entirely by a change in the plastic response to poor diet; on standard diet Selected and Control larvae showed nearly identical levels of all three BCAA. This suggests that selection driven by undernutrition favored a mechanism that resulted in poor diet-specific reduction in their abundance. This could have been because a low levels of BCAA in itself promote growth and development on poor diet, or because an increase in the rate of protein synthesis evolved by the Selected populations depletes these amino acids to a greater degree than most others.

Our results support the latter scenario. We showed that supplementing poor diet with BCAA improves growth of both Selected and Control larvae whereas adding three other essential amino acids does not, suggesting that one or more BCAA are limiting for growth. Leucine is the most abundant amino acid in the fly proteome (9.4% of predicted amino acids) with valine (6.2%) and isoleucine (5.2%) being in the second and sixth by abundance among essential amino acids, respectively (Piper *et al*. 2017). More importantly, while the latter two BCAA form a similar percentages of amino acids in yeast, leucine content in yeast protein is only about 7.5% (Piper *et al*. 2017). Thus, being in short supply proportionally to demand, leucine would be depleted more than other essential amino acids if the rate of protein synthesis increased, and thus might become the limiting amino acid. However, Piper *et al*. (2017) identified methionine as even more limiting; yet, in our study methionine is the only amino acid that tended to increase in Selected larvae relative to Control. Furthermore, the mismatch of amino acid composition in yeast and *Drosophila* proteins would not predict isoleucine and valine showing a similar decrease as a result of a higher protein synthesis rate. Thus, our results do not allow us to completely exclude the possibility that the depletion of BCAA in Selected larvae on poor diet is (also) mediated by reduced absorption from the diet and/or enhanced degradation. Deamination of all three BCAA is catalyzed by the same enzyme (Blumrich *et al*. 2021) while their intestinal absorption, at least in mammals, involves the same transmembrane transporters (Bröer & Fairweather 2018), which would explain parallel changes in abundance of all BCAA.

BCAA, and in particular leucine, play an important signaling role by stimulating the TOR pathway, which regulates protein synthesis (Antikainen *et al*. 2017). We have thus speculated that a reduction in the level of BCAA under nutrient shortage may have been favored by selection to avoid a situation where TOR signaling promotes initiation of more protein synthesis processes than the cells have amino acids to complete. However, if so, BCAA supplementation should have been harmful, which was not the case. Nonetheless, irrespective of the mechanism, the depletion of BCAA would have consequences for nutrient sensing, possibly requiring compensatory changes in growth regulation. This is consistent with changes in insulin signaling and the expression of FOXO targets we reported before (Erkosar *et al*. 2017; Kawecki *et al*. 2021). This interpretation of our results opens the question of potential evolutionary tradeoffs in nutritional adaptation that may result from the same compounds playing a role as regulatory molecules and building blocks of macromolecules or intermediates in energy metabolism.

## Supporting information

Raw_data_metabolomics

Raw_data_BCAA and WHK supplementation

Raw_data_15N enrichment

## Authors’ Contributions

FC and TJK designed the research, FC and LS performed all *Drosophila* experiments and collected sample, JES measured δ^15^N, HG-A and JI performed and oversaw metabolite quantification, FC, HG-A and TJK analyzed the data. FC and TJK wrote the paper, with contributions by all authors.

## Acknowledgements

This work has been supported by the Swiss National Science Foundation (grants number 31003A_162732 and 310030_184791 to TJK) and research funding from the University of Lausanne. We thank Nora Corthésy for help with the experiments and Berra Erkosar for her advice. The authors declare no conflicts of interest.

## Data Deposition

The metabolite abundance data, the data of larvae raised on BCAA or WHK or poor diet and the isotope content data are provided in supplemental data, respectively called “Raw_data_metabolomics”, “Raw_data_BCAA and WHK supplementation”, “Raw_data_15N enrichment”.

