## Supplementary Materials for "Experimental evolution of metabolism under nutrient restriction: enhanced amino acid catabolism and a key role of branched-chain amino acids"

#### Electronic Supplementary Material

| | |
|---|---|
| Detailed Methods | p. 2 |
| Supplementary Tables S1-S8 | p. 10 |
| Supplementary Figures S1-S6 | p.13 |
| Supplementary Data | separate files (“Raw_data_metabolomics”, “Raw_data_BCAA and WKK supplementation”, “Raw_data_15N enrichment”) |

### Detailed Methods

#### Experimental evolution

We used six Control and six Selected populations of *Drosophila melanogaster* originally derived from a sample of wild flies collected in Basel, Switzerland, in 1999 and adapted to the lab conditions over five years prior to the commencement of the evolution experiment. Details of the experimental evolution protocol are described in Kolss *et al.* (2009). Briefly, the Control populations have been maintained on 3-week generation cycle on a standard larval diet medium (15 g agar, 30 g sucrose, 60 g glucose, 12.5 g dry yeast, 50 g cornmeal, 0.5 g $CaCl_2$, 0.5 g $MgSO_4$, 10 mL Nipagin 10%, 6 mL propionic acid, 20 mL ethanol per liter of water), at a controlled density of approximately 200 eggs for 40 mL of food. The six Selected populations have been maintained with the same generation cycle and larval density on a poor larval diet containing one-fourth of sugars, yeast and cornmeal content present in the standard diet. After their emergence, adult flies of both regimes were transferred to standard diet and additionally fed with live yeast *ad libitum* before egg collection. Flies were reared at 25°C with 50–70% humidity and 12:12 light cycle. The experiments described in this paper were performed after 244 to 287 generations of experimental evolution.

Before each experiment, we reared the 12 populations for at least two generations on standard diet to minimize environmental maternal effects. To obtain larvae for each experiment, we let approximately 200 adult flies from each population lay eggs overnight on orange juice-agar plates supplemented with yeast. The desired number of eggs was then transferred onto the experimental media (see below) and inoculated with feces suspension from a pool of adult flies from all populations to ensure homogeneity of larval microbiota.

To prepare the feces inoculum we haphazardly collected 10-15 flies per population and transferred them into a petri dish containing a small slice of standard diet medium. After 48 h we removed the flies and the medium slice from the petri dish, washed the petri dish with PBS using a brush, filtered the resulting feces suspension through a fine mesh to remove eggs and debris, and adjusted the suspension to $OD_{600}$ = 0.5 with PBS. Each larval culture bottle was inoculated with 300 µL of this inoculum prepared on the same morning.

**High-coverage targeted metabolomics**

We performed high-coverage or multiple pathway targeted metabolomics, targeting several hundred metabolites involved in glycolysis, pentose phosphate pathway, purine and pyrimidine metabolism, TCA cycle, amino acid metabolism and oxidative phosphorylation. We used a factorial design whereby larvae from all Selected and Control populations were assayed on both diets, with three replicate samples per population and diet, in three batches per diet (one sample per batch and population). Due to workload samples from different diets were collected on different dates.

*Sampling*

In late third instar larvae a pulse of ecdysone is produced to initiate metamorphosis, which can have a large impact on metabolism and generate artifacts in the analysis (Lavrynenko *et al.* 2015; Li & Tennessen 2018; Kannangara *et al.* 2021). We aimed to study larval metabolism during growth phase before these pre-metamorphosis changes take place, in early third instar larvae. To assure a relative uniformity of their stage despite large differences in the rate of development due to diet and adaptation to it, we first transferred late second instar larvae (identified by "clubbed" anterior spiracles, http://www.shingletonlab.org/instars) from poor or standard diet to another dish of the same diet. Approximately 15 h later we collected samples of 15-20 larvae that transitioned to the third instar (recognizable by the presence of "branched" anterior spiracles). Following recommendations (Cox *et al.* 2017; Li & Tennessen 2018), larval samples were washed three times in 200 µL of PBS, then centrifuged at 2000 g and 4°C for 1 min to pellet larvae and remove any extra liquid that might interfere with mass spectrometry measurement. The pellets were snap-frozen and stored at -80°C.

*Multiple Pathway Targeted Analysis*

The samples were analyzed by Hydrophilic Interaction Liquid Chromatography coupled to tandem mass spectrometry (HILIC - MS/MS) in both positive and negative ionization modes using a 6495 triple quadrupole system (QqQ) interfaced with 1290 UHPLC system (Agilent Technologies). Data were acquired using two complementary chromatographic separations in dynamic multiple reaction monitoring mode (dMRM) as previously described (van der Velpen *et al.* 2019; Medina *et al.* 2020). Data were processed using MassHunter Quantitative Analysis (for QqQ, version B.07.01/Build 7.1.524.0, Agilent Technologies). Pooled QC samples were used to correct for signal intensity drift; metabolites with CV > 30% were discarded (Dunn *et al.* 2011; Tsugawa *et al.* 2014; Broadhurst *et al.* 2018). Linearity of metabolite response was verified with serially diluted quality controls (dQC); peaks for which the correlation with dilution factor was poor ($R^2 < 0.7$) were filtered out; resulting in 174 retained metabolites.

*Data transformation and outlier removal*

The raw values of peak areas were $log_2$ transformed, zero-centered and Pareto-scaled (i.e. divided by the square root of the standard error). Then, we identified outliers with a robust PCA on the covariance matrix of the data (Hubert *et al.* 2005). We used the R function *PcaHubert* with the number of principal components set to 2. Based on a visual inspection of the diagnostic plot, six outlier samples have been identified and removed (corresponding to three samples with a score distance above the cutoff and three samples with an orthogonal distance above 15; corresponding to four Selected samples, one from standard diet and three from poor diet and two Control samples reared on standard diet) (Supplementary Figure S6) (Hubert *et al.* 2005). We then fitted a linear mixed model for each metabolite, with the selection regime, the diet and their interaction as fixed factors and the replicate population, its interaction with diet and the batch as random factors. We then used *compute_redres* R function with "std cond" type to obtain Studentized residuals of each metabolite and sample. Thirty-two data metabolite-specific data points (belonging to 29 metabolites and 12 samples, 0.28% of all data point) with an absolute Studentized residual higher than 3.46 (likelihood of 0.001) were considered as outliers and removed. This last data set (i.e., 65 samples with 11278 data points) was used in the analyses described below (the data were not rescaled after outlier removal).

*Multivariate analysis*

To evaluate whether larvae reared on two different diets and from different selection regimes can be differentiated based on their overall metabolome, we performed a principal component analysis (PCA). using covariance matrix on log-transformed and Pareto-scaled concentrations (R package mixOmics (Rohart *et al.* 2017)). To avoid conflating two levels of replication (population and sample) and to reduce noise due to batch effects, we first obtained a single estimate of each metabolite's abundance per population and diet. To do so, we fitted a linear mixed model to the abundance of each metabolite (log-transformed and Pareto-scaled), with the population, the diet and their interaction as fixed factors and the batch as random factor. The estimated marginal means for each population × diet combination were obtained from this model (using R package emmeans) and used as input for the PCA and clustering heat map. To test whether the diet and evolutionary regime affect the position in the PC space, we performed a MANOVA on scores obtained from the first three PCAs as response variables, selection regime, diet and their interaction as fixed factors. We also performed clustering analysis using *heatmap* R function on the same data as the PCA.

*Univariate analysis*

To identify metabolites that differ between selection regime and diet, we fitted a linear mixed model (LMM) with Type 3 F-tests for each metabolite with *lmer* R function (Kuznetsova *et al.* 2017) and the same set of variables as in previous section. We then performed a contrast analysis in which we compared Control and Selected populations for each diet with *emmeans* and *pairs* functions in R. The p-values were corrected for multiple comparison using Storey q-value approach (pFDR; Storey & Tibshirani 2003). To help interpretation, we back-scaled contrast estimates towards $\log_2$ fold change (difference between Selected and Control populations or between poor and standard diets) by multiplying contrast estimates by their respective pareto factors.

We also used data from all metabolites to explore the quantitative relationship between the plastic response and the evolutionary response. Biologically it would be the most interesting to look at the relationship of the plastic response of the Control lines (i.e., $X_{\mathrm{CTLpoor}} - X_{\mathrm{CTLstd}}$) and the evolved relationship measured on the poor diet ($X_{\mathrm{SELpoor}} - X_{\mathrm{CTLpoor}}$), where $X$ stand for the estimated means of metabolites for conditions indicated by the subscripts. However, both are functions of the metabolite measurements in Control on poor diet. Consequently, the estimated covariance between them is biased by a component equal to $-\sigma^2_{\mathrm{CTLpoor}}$, where $\sigma^2$ stands for the error variance of $X$; this might produce an artifactual negative correlation in the absence of a real relationship. Therefore, we looked instead at the correlation between the main effects of the diet (corresponding to $X_{\mathrm{SELpoor}} + X_{\mathrm{CTLpoor}} - X_{\mathrm{SELstd}} - X_{\mathrm{CTLstd}}$) and the main effect of evolutionary regime (corresponding to $X_{\mathrm{SELpoor}} - X_{\mathrm{CTLpoor}} + X_{\mathrm{SELstd}} - X_{\mathrm{CTLstd}}$). While the two composite variables are functions of the same measurements, their error covariance is $(\sigma^2_{\mathrm{SELpoor}} - \sigma^2_{\mathrm{CTLpoor}} - \sigma^2_{\mathrm{SELstd}} + \sigma^2_{\mathrm{CTLstd}})/4$; it vanishes if the error variances of the different conditions are the same.

Finally, we tested whether Selected and Control populations differ in overall plastic response of metabolome to diet populations by fitting a major axis regression to $X_{\mathrm{CTLpoor}} - X_{\mathrm{CTLstd}}$ and $X_{\mathrm{SELpoor}} - X_{\mathrm{SELstd}}$ (R package smatr (Warton *et al.* 2012)). A systematic difference in the magnitude of response would be detected as a deviation of the major axis slope from 1.

**Developmental rate and survival in food enriched with amino acids**

In this experiment the larvae of all Selected and Control populations were raised on three diet treatments: the poor diet (same as in experimental evolution), poor diet supplemented with the three branched chain amino acids (BCAA) and poor diet supplemented with three other essential amino acids: tryptophan (W), histidine (H) and lysine (K). The concentration of these amino acids in the poor diet was estimated based on their content in yeast (Piper *et al.* 2017), and we aimed to double their concentrations in the respective treatments. To this end, the BCAA-supplemented diet consisted of the poor diet supplemented with 1.5 mM valine, 1.8 mM leucine and 1.3 mM isoleucine (Sigma-Aldrich, Switzerland). Similarly, the WHK-enriched diet was supplemented with 0.5 mM, 0.8 mM and 1.9 mM of tryptophan, histidine and lysine, respectively.

Each replicate bottle with 40 mL of one of these food media was seeded with 200 eggs and inoculated with feces as described above. Upon emergence, adult males and females were removed and counted once a day until day 26 counted from egg-laying. These data were used to estimate for each replicate egg-to-adult survival probability and the mean developmental rate (mean of the inverse developmental time). From each bottle 10 females were randomly collected on the day of peak emergence (i.e., when the largest number of females emerged). These female samples were frozen and dried at 70°C for 24h before to be weighed as a group to the nearest 1 µg on a precision balance (Mettler Toledo, MT5); sample weight was divided by the number of females weighed (i.e., by 10, or 9 in replicates where a female escaped) to obtain female weight. We also estimated the average growth rate of the larvae in each bottle as ln(female weight/egg weight)/larval developmental time, where egg weight was assumed to be 5 µg (Kolss *et al.* 2009) and the larval developmental time was estimated as the time from egg laying to the day on which the females were collected minus 5 days to account for the time needed for egg development and metamorphosis. This experiment was done in three blocks with one replicate per population, diet treatment and block.

To test for the effect of diet treatment and evolutionary regime on developmental rate, female dry weight and growth rate, we fitted a linear mixed models using lmer package of R with the evolutionary regime, treatment, sex (for developmental rate only) and their interactions as fixed factors, the replicate population, block and bottle (for developmental rate only) as random factors. The egg counting and sorting was done by two experimenters with three Control and three Selected populations each (the populations-experimenter pairs were kept the same along the three blocks). Because the egg manipulation could affect the eclosion rate, we included experimenter as a random factor in the models mentioned above. The effects were tested with F tests; the degrees of freedom were estimated with the Satterthwaite approximation. For survival, we fitted a generalized mixed model with binomial error distribution and logit link with the number of emerged versus non emerged flies as the response, with the same fixed and random factors as for growth rate. Significance of the fixed effects was tested with likelihood ratio tests using afex package (Singmann *et al.* 2015); pairwise and custom contrasts were estimated and tested with emmeans package in R.

**Stable isotope experiment**

This experiment aimed to compare accumulation of the heavy stable isotope of nitrogen ($^{15}N$) as an indicator of the proportion of absorbed amino acids that were subject to deamination followed by excretion of nitrogen as waste. Amino acid deamination process has a slight preference for the common light nitrogen isotope ($^{14}N$); as a consequence, proteins in tissues become enriched in $^{15}N$ to a degree that increases with rate of amino-acids deamination (Minagawa & Wada 1984; Peterson & Fry 1987; Gannes *et al.* 1997). Because $^{15}N$ is naturally rare, in this experiment we used food media containing yeast enriched in $^{15}N$, which improved signal to noise ratio.

*Food media preparation*

In this experiment we used modified versions of standard and poor media that differed from those used in the course of experimental evolution and in the other experiments described above in two ways (Cavigliasso *et al.* 2020). First, they did not contain cornmeal, leaving yeast as the only source of nitrogen. Second, we used $^{15}N$-enriched baker's yeast in addition to dead brewer's yeast used for the usual standard and poor diets. We grew the baker's yeast (starting from a dry form, $OD_{600}$=100) overnight at 30°C in $^{15}N$-enriched SD media (8.5 g yeast nitrogen base without amino acids and ammonium sulfate, 2.5 g normal ammonium sulfate, 2.5 g $^{15}N$-labeled ammonium sulfate ($(^{15}NH_4)_2SO_4$ and 20 g sucrose per liter of water). This $^{15}N$-enriched yeast was collected by removing the supernatant after centrifugation (5 min at 3000g) and stored at -20°C. To prepare "$^{15}N$-enriched standard diet", we added 2.5 mL of $^{15}N$-enriched cultured yeast to a suspension of 12.5 g of dry baker's yeast in 100 mL of water before transferring it in one liter of water solution containing 10 g agar, 90 g of sucrose, 0.5 g $CaCl_2$, 0.5 g $MgSO_4$, 10 mL Nipagin 10%, 6 mL propionic acid and 20 mL ethanol. After solidification, we blended the diet medium and transferred to petri dishes (10 mL per dish of 46 mm diameter). The corresponding "$^{15}N$-enriched poor diet" was prepared in the analogous way, but one-fourth of the amounts of sugar and yeast of both kinds were used.

*Sample collection*

To raise individuals for isotope measurements we transferred approximately 150 eggs to petri dishes with $^{15}N$-enriched standard or poor diet. For each sample, we collected 20-30 prepupae and 20-30 virgin adult females, pooled from two (standard diet) or three (poor diet) petri dishes; the females were starved for three hours on agar to permit emptying of their guts. The samples were flash-frozen and stored at -80°C. Two samples were collected per population and diet, in two blocks per diet carried out on different dates. Because the level of $^{15}N$ in diet may differ between block and diet and thus may affect the level in tissue sample, we also collected a "diet sample" for each diet and block.

*Isotope measurement*

We followed identical protocol to (Cavigliasso *et al.* 2020): The relative $^{15}N$ content of the samples was determined by elemental analysis/isotope ratio mass spectrometry (EA/IRMS), using a Carlo Erba 1108 (Fisons Instruments, Milan, Italy) elemental analyzer connected to a Delta V Plus isotope ratio mass spectrometer via a ConFlo III split interface (both of Thermo Fisher Scientific, Bremen, Germany), operated under continuous helium (He) flow (Spangenberg *et al.* 2006; Spangenberg *et al.* 2020). The nitrogen stable isotope compositions were reported in the delta ($\delta$) notation as per mil (‰) variations of the molar ratio ($R$) of the heavy ($^{15}N$) to light isotope ($^{14}N$) of nitrogen relative to the international standard molecular nitrogen in air ($N_2$-Air).

$$\delta^{15}\mathrm{N}_{\mathrm{sample/standard}} = \frac{R\left(\frac{15\mathrm{N}}{14\mathrm{N}}\right) sample}{R\left(\frac{15\mathrm{N}}{14\mathrm{N}}\right)\mathrm{N2-Air}} - 1$$

For calibration and normalization of the measured $\delta^{15}N$ values to $N_2$-Air scale, a 3- point calibration was used with international reference materials (RMs) and University of Lausanne in-house standards (Spangenberg *et al.* 2020). The nitrogen concentration (in wt.%) were determined from the peak areas of the major isotopes using the calibrations for $\delta^{15}N_{Air-N2}$. The repeatability was better than 0.2 wt.%.

*Statistical analysis*

To account for potential variation of $\delta^{15}N$ values between diet types and blocks, $\delta^{15}N$ values in prepupae and adults were normalized by subtracting their $\delta^{15}N$ vales from the corresponding diet and block. These normalized values ($\Delta^{15}N$) were used as the response variable. We fitted a linear mixed model (LMM) with Type 3 F-tests with the selection regime, developmental stage, diet and their interactions as fixed factors, the replicate populations, its interaction with stage and blocks as random factors.

#### Supplementary Tables

**Supplementary Table S1.** Summary of results of the ANOVAs on scores from the first three principal components of metabolite abundance.

| Factor | PC1 | | PC2 | | PC3 | |
|---|---|---|---|---|---|---|
| | Statistics | *P* | Statistics | *P* | Statistics | *P* |
| Regime | $F_1 = 0.1$ | 0.71 | $F_1 = 14.3$ | **0.001** | $F_1 = 13.7$ | **0.001** |
| Diet | $F_1 = 1274.7$ | **< 0.001** | $F_1 = 0.2$ | 0.68 | $F_1 = 0.1$ | 0.82 |
| Regime × Diet | $F_1 = 2.1$ | 0.17 | $F_1 = 1.7$ | 0.21 | $F_1 = 1.2$ | 0.29 |

**Supplementary Table S2.** Summary of univariate analysis with q-values and log2 fold change. A separate file.

**Supplementary Table S3.** Significance tests from linear mixed models (LMM) for the four amino acids that show significant diet × regime interaction: BCAA (leucine, isoleucine and valine) and proline, reported in Figure 4A-D. SEL: Selected populations; CTL: Control populations; Std: standard diet. *q* are the corrected *P* (10% FDR).

| Factors | Leucine | | Isoleucine | | Valine | | Proline | |
|---|---|---|---|---|---|---|---|---|
| | Statistics | *q* | Statistics | *q* | Statistics | *q* | Statistics | *q* |
| Regime | $F_{1,13} = 6.9$ | **0.061** | $F_{1,12} = 8.6$ | **0.044** | $F_{1,12} = 5.8$ | **0.073** | $F_{1,12} = 1.1$ | 0.31 |
| Diet | $F_{1,54} = 0.2$ | 0.70 | $F_{1,5} = 2.6$ | 0.17 | $F_{1,6} = 7.2$ | **0.038** | $F_{1,4} = 10.2$ | **0.031** |
| Regime × Diet | $F_{1,54} = 14.3$ | **0.018** | $F_{1,48} = 16.7$ | **0.011** | $F_{1,49} = 18.4$ | **0.011** | $F_{1,12} = 13.8$ | **0.071** |
| *Pairwise Contrasts* | | | | | | | | |
| SEL – CTL in poor diet | $t_{25} = 4.1$ | **0.004** | $t_{24} = 4.4$ | **0.003** | $t_{24} = 4.4$ | **0.003** | $t_{20} = 2.3$ | **0.073** |
| SEL – CTL in std diet | $t_{24} = 0.3$ | 0.77 | $t_{23} = 0.0$ | 0.96 | $t_{23} = 0.9$ | 0.56 | $t_{20} = 0.5$ | 0.62 |
| Poor – Std in SEL | $t_6 = 2.8$ | **0.051** | $t_{12} = 3.5$ | **0.014** | $t_{13} = 4.1$ | **0.006** | $t_{14} = 0.1$ | 0.95 |
| Poor – Std in CTL | $t_6 = 2.3$ | **0.057** | $t_{11} = 1.4$ | 0.19 | $t_{13} = 0.3$ | 0.79 | $t_{14} = 4.2$ | **0.002** |

**Supplementary Table S4**: Amino acid supplementation experiment: summary of significance tests from generalized linear mixed model for egg-to-adult survival rate reported in Figure 4E. Treatment refers to the three types of diet treatment (poor diet, BCAA supplementation, WHK supplementation).

| Factors | Statistics | *P* |
|---|---|---|
| Regime | $\chi^2 = 1.5$ | 0.21 |
| Treatment | $\chi^2 = 3.4$ | 0.18 |
| Regime × Treatment | $\chi^2 = 0.1$ | 0.97 |

**Supplementary Table S5.** Amino acid supplementation experiment: summary of significance tests from LMM for developmental rate reported in Figure 4F. Treatment refers to the three types of diet treatment (poor diet, BCAA supplementation, WHK supplementation).

| Factors | Statistics | *P* |
|---|---|---|
| Regime | $F_{1,9} = 245.5$ | < 0.001 |
| Treatment | $F_{2,91} = 36.8$ | < 0.001 |
| Sex | $F_{1,10} = 3.9$ | 0.075 |
| Regime × Treatment | $F_{2,91} = 4.2$ | 0.018 |
| Regime × Sex | $F_{1,10} = 1.0$ | 0.34 |
| Treatment × Sex | $F_{2,22} = 1.2$ | 0.32 |
| Regime × Treatment × Sex | $F_{2,22} = 3.6$ | 0.046 |
| *Pairwise Contrasts* | | |
| For Control | | |
| BCAA – Poor | $t_{21} = 3.5$ | 0.006 |
| WHK – Poor | $t_{20} = -0.4$ | 0.90 |
| BCAA – WHK | $t_{21} = 4.0$ | 0.002 |
| For Selected | | |
| BCAA – Poor | $t_{20} = 4.6$ | < 0.001 |
| WHK – Poor | $t_{20} = 3.4$ | 0.008 |
| BCAA – WHK | $t_{20} = 8.0$ | < 0.001 |
| *Custom Interaction Contrasts* | | |
| $(BCAA – Poor)_{in\ SEL}$ – $(BCAA – Poor)_{in\ CTL}$ | $t_{20} = 0.7$ | 0.48 |
| $(WHK – Poor)_{in\ SEL}$ – $(WHK – Poor)_{in\ CTL}$ | $t_{20} = -2.1$ | 0.051 |

**Supplementary Table S6**: Amino acid supplementation experiment: summary of significance tests from LMM for female weight reported in Figure 4G. Treatment refers to the three types of diet treatment (poor diet, BCAA supplementation, WHK supplementation).

| Factors | Statistics | *P* |
|---|---|---|
| Regime | $F_{1,10} = 88.3$ | < 0.001 |
| Treatment | $F_{2,20} = 2.0$ | 0.16 |
| Regime × Treatment | $F_{2,20} = 1.8$ | 0.19 |

**Supplementary Table S7.** Amino acid supplementation experiment: summary of significance tests from LMM for female growth rate reported in Figure 4H. Treatment refers to the three types of diet treatment (poor diet, BCAA supplementation, WHK supplementation).

| Factors | Statistics | *P* |
|---|---|---|
| Regime | $F_{1,9} = 21.5$ | 0.001 |
| Treatment | $F_{2,20} = 9.0$ | 0.002 |
| Regime × Treatment | $F_{2,20} = 1.4$ | 0.26 |
| *Pairwise comparisons* | | |
| BCAA – Poor | $t_{20} = 2.8$ | 0.029 |
| WHK – Poor | $t_{20} = -1.4$ | 0.36 |
| BCAA – WHK | $t_{20} = 4.2$ | 0.001 |

**Supplementary Table S8.** Summary of type 3 F-tests with Satterthwaite's method for fixed effect in LMM on $\Delta^{15}N$ (‰ $N_2$-Air) reported in Figure 5C.

| Factors | F value | *P* |
|---|---|---|
| Regime | $F_{1,11} = 18.9$ | 0.001 |
| Stage | $F_{1,71} = 18.4$ | <0.001 |
| Diet | $F_{1,12} = 88.4$ | <0.001 |
| Regime × Stage | $F_{1,71} = 1.5$ | 0.22 |
| Regime × Diet | $F_{1,12} = 2.8$ | 0.12 |
| Diet × Stage | $F_{1,71} = 0.9$ | 0.34 |
| Regime × Stage × Diet | $F_{1,71} = 0.2$ | 0.63 |

#### Supplementary Figures

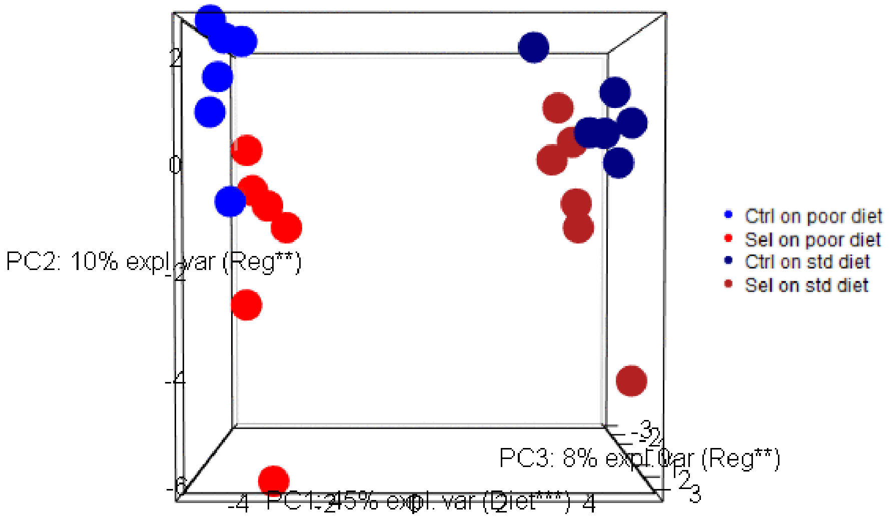


**Supplementary Figure S1.** 3D animation of sample clustering using the first 3 principal components of multiparametric metabolic profiles composed of 174 metabolite abundance data; each point corresponds to a population estimate averaged over replicate samples.

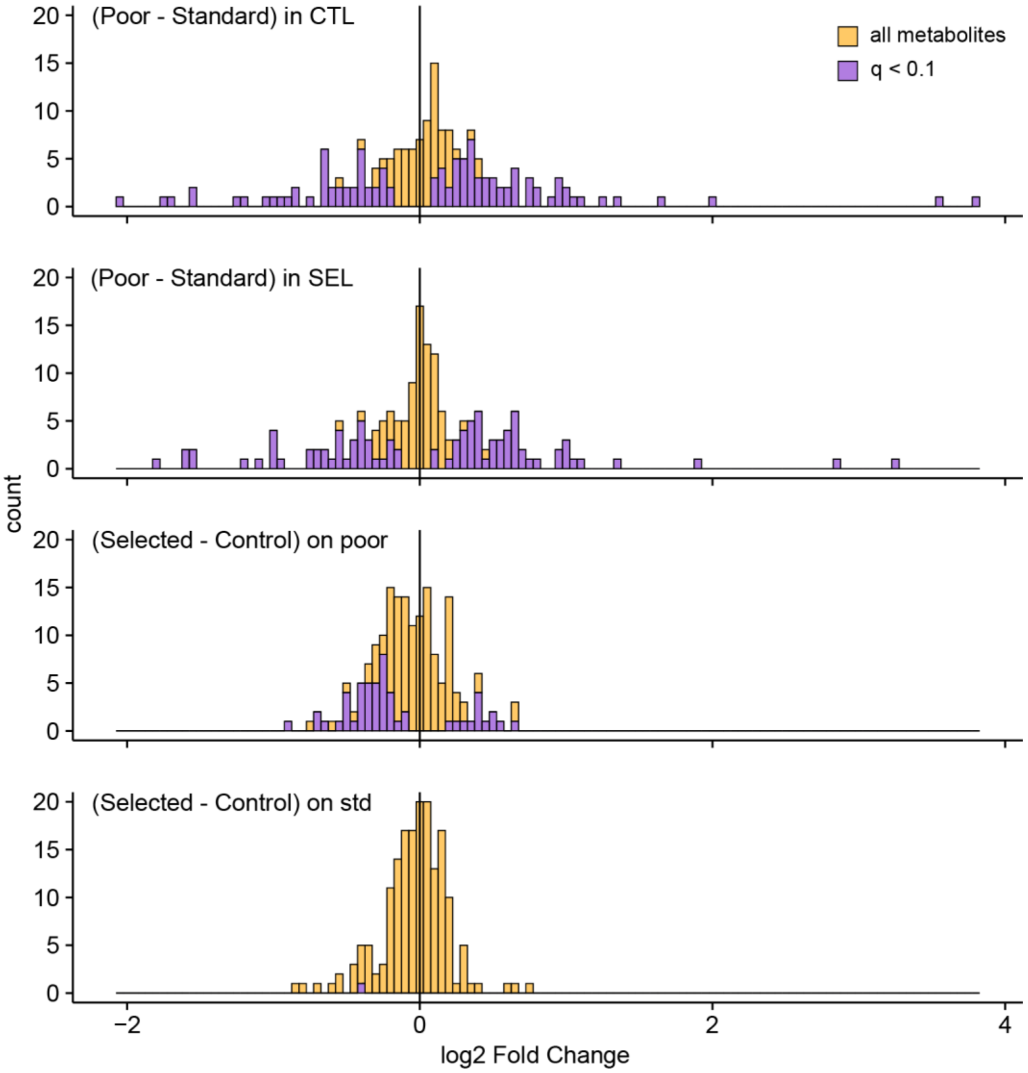


**Supplementary Figure S2:** Distribution of differences in metabolite abundance for different pairwise contrasts. CTL: Control; SEL: Selected; poor: poor diet; std: standard diet. Purple bars: metabolites significant ($q < 0.1$) for the particular contrast.

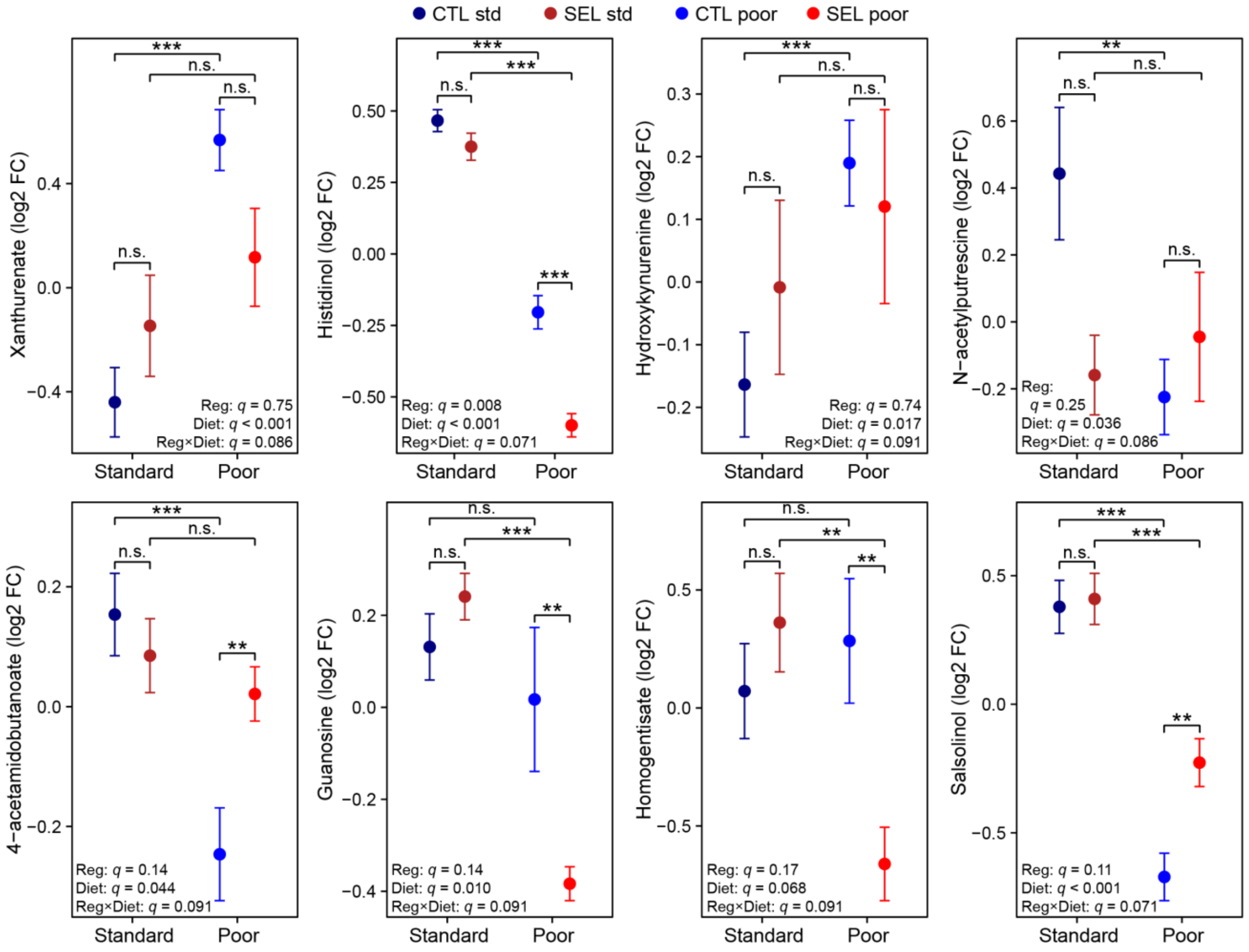


**Supplementary Figure S3.** Relative abundance of metabolites with significant regime × diet interaction ($q < 0.1$) in Selected (SEL) and Control (CTL) larvae raised on poor and standard (std) diet. Level of significance: n.s not significant $q > 0.1$; * $q < 0.1$; ** $q < 0.05$; *** $q < 0.01$. Symbols indicate means ± SE. N = 3 batches (pool of 15-20 larvae) per population and diet. Reg: evolutionary regime.

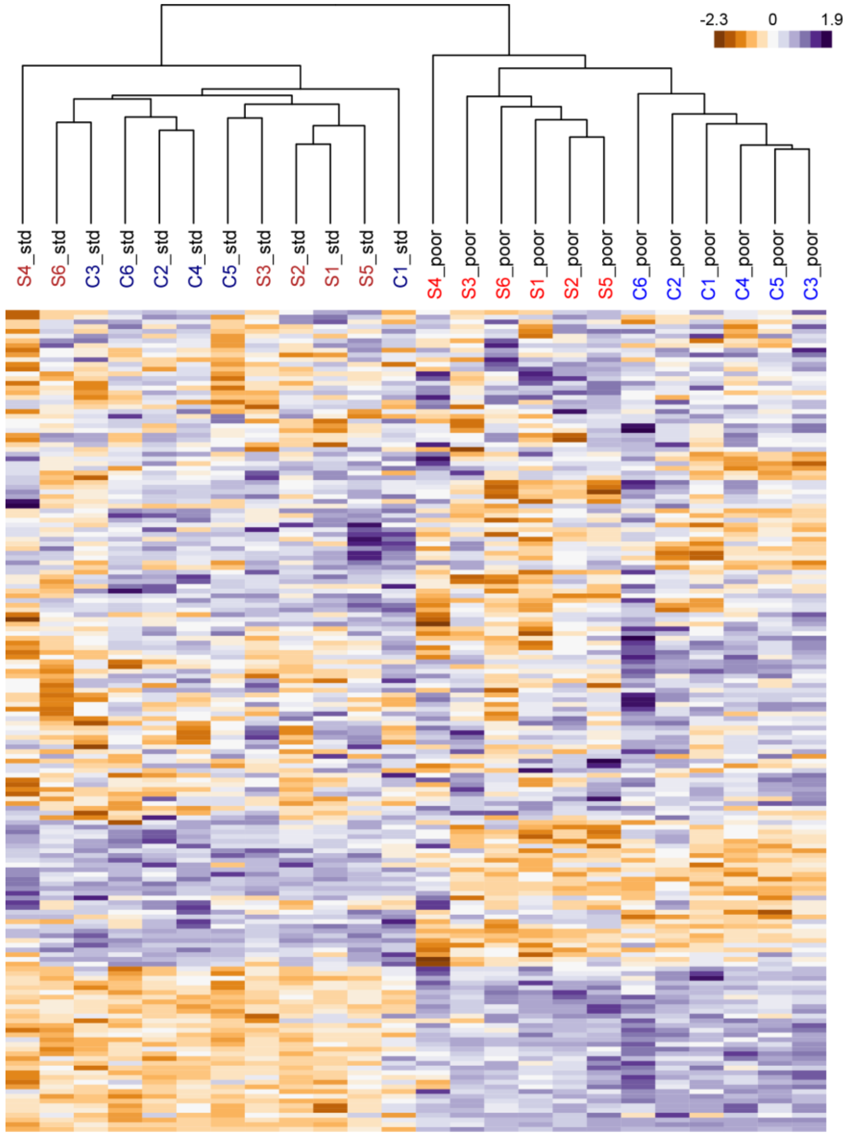


**Supplementary Figure S4:** Clustering of populations based on their metabolic profiles. Each cell of the heat map is colored according to the metabolite abundance (estimate averaged over replicate samples, as for the PCA in Figure 1A). S1-S6: Selected populations, C1-C6: Control populations; "_poor": larvae raised on poor diet, "_std": larvae raised on standard diet.

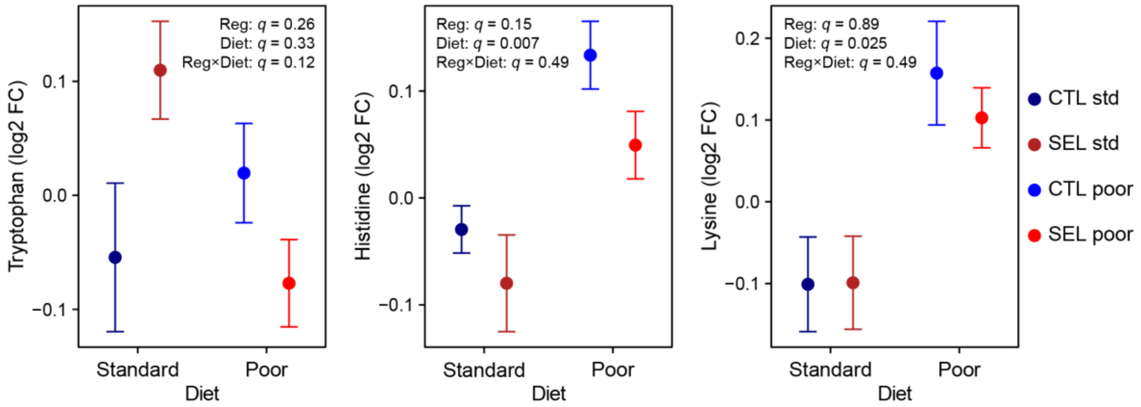


**Supplementary Figure S5.** Relative amount of tryptophan, histidine and lysine in Selected (SEL) and Control (CTL) larvae raised on poor and standard (std) diet. Symbols indicate means ± SE. N = 3 batches (pool of 15-20 larvae) per population and diet. Reg: evolutionary regime.

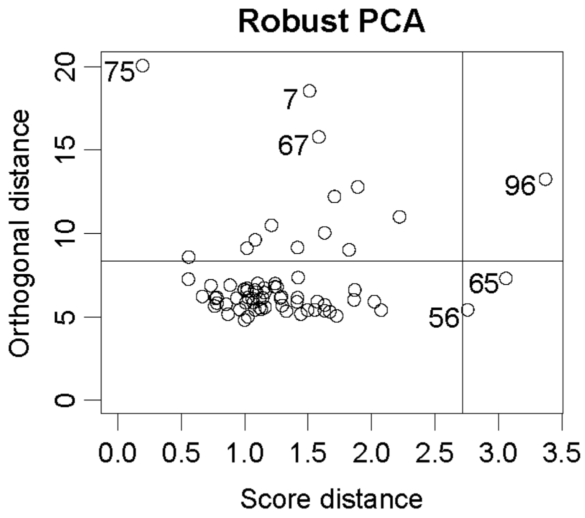


**Supplementary Figure S6.** Robust PCA fitted with two components. Based on visual inspection, six outlier samples (labeled with sample number) were removed for the analysis. The two lines show the conventionally assumed cutoffs (for details see (Hubert *et al.* 2005)).
