## Supplementary figures and images for "Experimental evolution of metabolism under nutrient restriction: enhanced amino acid catabolism and a key role of branched-chain amino acids"

### Supplementary Figure S1

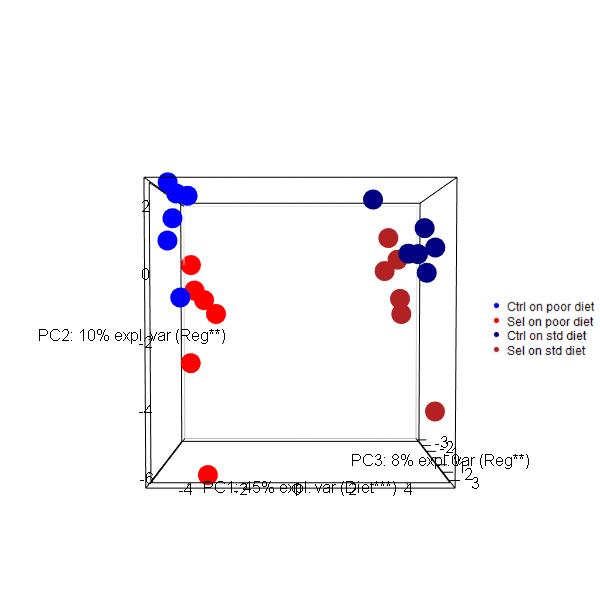
